# Investigating the folding dynamics of NS2B protein of Zika virus

**DOI:** 10.1101/2022.12.12.520106

**Authors:** Ankur Kumar, Prateek Kumar, Pushpendra Mani Mishra, Rajanish Giri

## Abstract

NS2B protein of the Zika virus acts as a co-factor for NS3 protease where only the cytosolic domain of NS2B is sufficient for the protease activity. At the same time, NS2B also involves in remodeling the NS3 protease structure. In isolation, we previously proved the NS2B cytosolic domain (residues 49-95) as a disordered type peptide conformation. Further, this study investigated the overall dynamics of NS2B full-length protein. Our Alphafold2 structure modeling system revealed surprising similarities between selected flavivirus NS2B proteins. This similarity reflects that the NS2B protein across flavivirus is conserved fold-wise. The MD simulation of Zika virus NS2B full-length protein shows that the cytosolic domain as a part of full-length protein is a disorder region supporting our previous experimental finding, which suggests the disordered nature of the cytosolic domain in isolation. Since the cytosolic domain of NS2B is essential for protease activity, we have also investigated the folding and dynamics of the NS2B cytosolic domain (residues 49-95) that shows the disorder to alpha helix transition in TFE. On the other hand, in the presence of SDS, macromolecular crowder like ficoll and PEG do not induce secondary structural change. This dynamics study could have implications for some unknown folds of the NS2B protein.

## 1. Introduction

The Zika virus (ZIKV) genome encodes a single polyprotein co-translationally incorporated in the endoplasmic reticulum (ER) [1] and cleaves into three non-structural and seven non-structural proteins. This polyprotein is hydrolyzed by the host-cell signalase and viral NS2B-NS3 protease at the lumen of ER and the cytoplasmic side, respectively. Subsequently, the host-cell furin cleaved off the pre-membrane (prM) in the trans-Golgi network [1]. Furthermore, in NS2B-NS3 protease, NS2B act as a co-factor required for the catalytic activity of NS3 protease [2].

Like other flaviviruses [3–5], the ZIKV NS2B is a membrane protein consisting of two transmembrane domains (C-terminal and N-terminal) and a cytosolic domain (residues 45-96) [6], where only the cytosolic domain is sufficient for the protease activity [1,7,8]. Furthermore, the cytosolic domain is hydrophilic, whereas the C-terminal and N-terminal domain is hydrophobic that tightly associated with ER membrane to provide a membrane anchor for NS3 protease [2,6,9]. Notably, a mutation in NS2B β-strand impedes the NS2B-NS3 interaction and reduces the ability of NS3 protease to hydrolyze the polyprotein [2]. Additionally, ZIKV NS2B forms an oligomer [10], like other flaviviruses, that might be essential for the viral replication [5]. Besides, NS2B in isolation mainly comprises helices and disordered regions in micelles determined by NMR [10].

Previously, in isolation, the cytosolic domain of the NS2B was predicted to have 37 residues long disordered secondary structure (residues 68-98) towards the C-terminus of this domain [11,12]. However, in complex with NS3 protease, the NS2B co-factor (residues 45-96) comprises four β-strand (51-57, 66-67, 73-78, and 84-86) [7,8]. The co-factor NS2B wraps NS3 protease so that the C-terminus (residues 71-87) of this protein forms a β-hairpin. This β-hairpin lies at the NS3 protease active site and interacts with the protease substrate at the active site. Further, residues 74 to 86 of this co-factor shape the protease S2 pocket [1,13,14]. Notably, the C-terminus part (residues 45-96) of NS2B stabilized at the active site by interacting with Ser 135, Gly 133, and His 51 of the protease, leading to the protease structure remodeling. This structural change enables the binding of an incoming substrate at the active site. Therefore this evidence suggests the dual role of NS2B (45-96) in NS3 protease activation by inducing protease refolding and interacting with the substrate [1,15].

The previous literature shows minimal understanding of ZIKV NS2B dynamics in isolation. Therefore, this study investigated the overall dynamics of NS2B full-length protein using molecular dynamics simulation studies. However, the primary role of NS2B as a co-factor, different regions of NS2B protein, viz. transmembrane domain, cytosolic domain, N-terminal tail, and C terminal tail, have different functional roles during the replication and maturation of ZIKV. So it is essential to investigate the dynamics of each segment separately. In this study, our further goal was to analyze the conformational dynamics of the cytosolic domain (residues 49-95) using different probes or solvent conditions. We used TFE as a potent alpha helix inducer to analyze the extent of dynamics and the conformational transition in NS2B cytosolic domain to check how many different conformations this domain can attain. Using a combination of theoretical (simulations) and experimental approaches (CD and Fluorescence), we deciphered the overall dynamics of NS2B protein and its cytosolic domain in isolation by employing a reductionist approach.

## 2. Material and methods

### 2.1. peptide and buffer

NS2B cytosolic domain peptide with >97 % purity (residues 49-95; VDMYIERAGDITWEKDAEVTGNSPRLDVALDESGDFSLVEDDGPPMA) was purchased from Thermo Fisher Scientific. The isoelectric point of this peptide is 3.73, determined by the Expasy Protparam tool. NS2B (residue 49-95) peptide was dissolved in 50 mM phosphate buffer, pH 7, to prepare a stock solution of 195 µM. The buffer condition of the protein sample was 50 mM Na-phosphate buffer, pH 7 for each experiment except for TFE (Milli-Q water was used instead of buffer).

### 2.2. Circular dichroism (CD) measurement

CD measurement of NS2B cytosolic domain (residue 49-95) was monitored using a CD spectrophotometer (J1500, Jasco). 10 μM of NS2B peptide was used to record the far UV-CD (190-240 nm; on an interval of 0.5 nm) spectra in a quartz cuvette (Hellma, Plainview, NY, USA) of 1mm pathlength at 25°C for an average of 3 consecutive scans. Each spectrum was subjected to smoothing by Savitsky-Golay methods by selecting 2^nd^ polynomial order and 20 data points.

### 2.3. Fluorescence measurement

The intrinsic fluorescence probe Tyr at the 61^st^ residue position in NS2B (residue 49-95) peptide was used to monitor fluorescence emission spectra using a Fluorolog spectrofluorometer (Horiba Scientific). Trp was excited at 295 nm wavelength and monitored the emission spectra between 300-550 nm with 5 nm slit bandwidth. In addition, 5 μM of NS2B (residue 49-95) was used to record the spectra at 20 °C.

### 2.4. Fluorescence lifetime measurement

The fluorescence lifetime of the intrinsic fluorescence probe, Trp in NS2B (residue 49-95) peptide, was monitored using DeltaFlex TCSPC system (Horiba Scientific). 5 µM peptide was used to monitor the spectra where the range of measurement was set to 200 ns with 10000 counts of peak preset and 32 nm bandpass. The instrument response factor (IRF) was corrected using ludox by measuring prompt scan at 284 nm wavelength. 350 nm wavelength was set for the emission monochromator. All the lifetime curve was fitted using the three-exponential decay function.

### 2.5. Structure modeling

The structures of NS2B full-length protein (130 residues long) and an isolated cytosolic domain (residues 49-95) were modeled using I-TASSER web server, as described previously[16] and AlphaFold2 modeling server[17]. The I-TASSER web server generated five model structures, and we selected the model with the highest confidence score (c-score) and cluster density. Similarly, the AlphaFold2 modeling system generated five model structures, where we selected the ranked_0 structure for further analysis. For structural optimization and minimization, the structure models were processed in protein preparation wizard in Schrodinger [18,19] using OPLS2005 forcefield [20,21].

### 2.6. Molecular dynamics (MD) simulation

The Gromacs simulation package was utilized for all simulations of the NS2B protein of ZIKV. The NS2B full-length protein comprises transmembrane-spanning regions at the N-terminal and C-terminal domains. Therefore, it was placed inside a bilayer membrane consisting of 150 molecules of DOPC lipids in both leaflets. The charge neutralization was done using Potassium and Chloride ions. For this purpose, the CHARMM-GUI server was utilized, and then the parameters of minimization and equilibration were obtained for Charmm36 forcefield to be executed in Gromacs. The minimization was carried out up to 5000 steps Using the steepest descent method. The equilibration step was performed for 500 ps with Berendsen pressure and temperature coupling algorithms. Finally, the production run was executed for 100 ns with the Nose-Hoover method of temperature coupling (average temperature 303.15 K) and Parrinello-Rahman method of pressure coupling (average pressure 1 bar). For hydrogen-bond constraints, LINCS algorithm was applied.

Since we have chosen an isolated NS2B cytosolic domain (residues 49-95), we performed MD simulations for NS2B cytosolic domain (residues 49-95) without any lipid membranes. First, we performed simulations of the modeled structure to obtain an energy-minimized structure. Therefore 500 ns (NS2B_49-95_ model structure predicted by I-TASSER) and 100 ns (NS2B_49-95_ model structure predicted by AlphaFold2) long simulations were performed in aqueous conditions containing SPC water model with charge neutralization by Sodium ions. Then, the last frame of NS2B (residues 49-95) model structure (I-TASSER) trajectory was used for further simulations in different solvent conditions, such as TFE up to one microsecond. For TFE, the topology parameters were generated, and the subsequent forcefield parameters for GROMOS54a7 were obtained using PRODRG server [22]. For each simulation setup, 50000 steps were run for minimization using the steepest-descent method. After minimization, the equilibration process with NVT and NPT ensembles for 100 ps each was carried out with modified Berendsen or V-rescale, a temperature coupling method, and a Parrinello-Rahman algorithm for pressure coupling. Finally, our in-house computing facility performed an extensively long MD production run for each simulation setup up to 1 μs at 300 K temperature and 1 bar pressure.

## 3. Results

### 3.1. Structure modeling of NS2B full-length protein of ZIKV

We used the amino acid sequence from the Brazilian isolate of ZIKV (BeH823339) NS2B full-length (130 residues long) protein for the structural modeling. This protein sequence was subjected to modeling using AlphaFold2 and I-TASSER modeling system. The ranked_0 structure in the AlfaFold2 modeling system was selected as a template structure among the top five (Figure 1A). The template structure was observed to have four α-helix (6-23, 30-44, 95-109, and 114-128) at the N-terminal (1-44) and C-terminal domain (96-130) and two β-strands (76-78 & 84-86) at the cytosolic domain (45-96).

**Figure 1.**
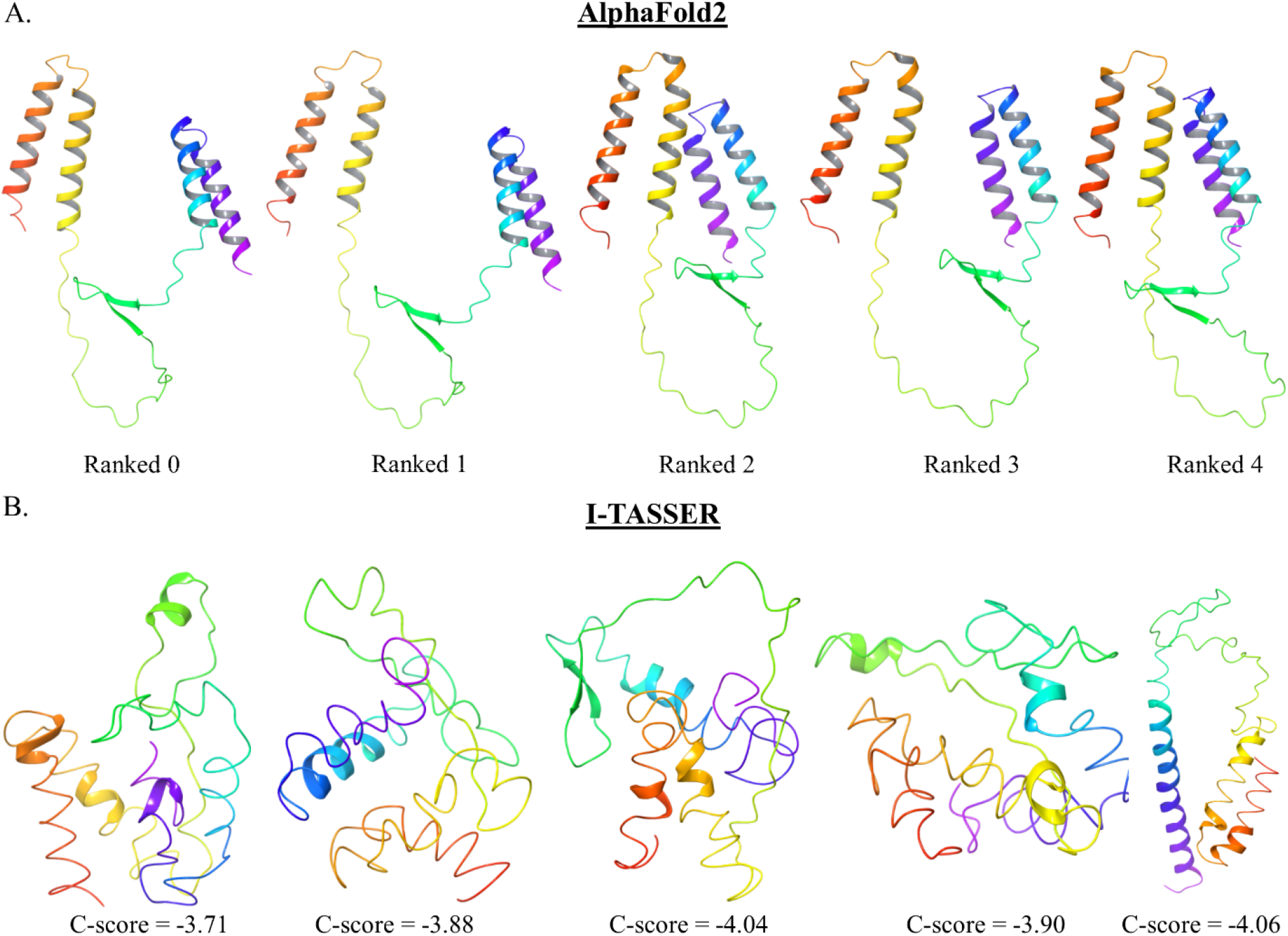
Structure modeling of ZIKV NS2B full-length protein. (A) model structures of ZIKV NS2B full-length (130 residues long) protein from AlphaFold2 modeling system. The ranked_0 structure is used as a template to analyze the structural elements. The red-colored terminal end represents the N-terminal (1^st^ residue), and the purple-colored terminal end represents the C-terminal (130^th^ residue). (B) Model structures of ZIKV NS2B full-length (130 residues long) protein from I-TASSER modeling system. The structure with a C-score of - 3.71 is the lowest energy model used to analyze the structural elements. The red-colored end is the N-terminal (1^st^ residue), and the purple-colored end is the C-terminal (130^th^ residue).

Additionally, we predicted the ZIKV NS2B full-length structure using I-TASSER modeling system, as shown in Figure 1B. The structure with a confidence score (C-score) of - 3.71 wa used as a template structure. The template structure consists of two α-helix (18-21, 33-36) at the N-terminal, one α-helix (119-122) at the C-terminal domain, and one α-helix (63-67) at the cytosolic domain (49-95). In both the prediction system, the cytosolic domain (49-95) of NS2B protein is showing high content of disordered structure, in line with our previously determined NS2B cytosolic domain (49-95) structure as a disordered type protein structure (determined by disordered prediction server and the CD spectroscopy) [11].

Furthermore, we did a comparative analysis of the sequence of ZIKV NS2B full-length protein with a few members of flaviviruses like DENV, JEV, WNV, YFV, TBEV, MVEV, OHFV, and St. Louis encephalitis virus (St. Louis EV) using Clustal omega multiple sequence alignment software (Figure S1). As a result, ZIKV NS2B full-length protein shows ∼24-48 % sequenc identity with these members (Table 1). Additionally, we prepared a structure model of NS2B protein of these members by AlfaFold modeling system to compare the secondary structure w.r.t ZIKV (Figures 2, 3 & 4).

**Table 1.** A summary of the secondary structural elements in NS2B protein (modeled from AlphaFold2) of the few members of flaviviruses.

| Virus | helix |  |  | strand |  | helix |  | % identity with ZIKV | RMSD (Å) w.r.t ZIKV |
| --- | --- | --- | --- | --- | --- | --- | --- | --- | --- |
| ZIKV | 6-22 | 29-47 | - | 76-78 | 84-86 | 95-109 | 114-128 | - | - |
| DENV1 | 5-21 | 30-46 | - | 76-78 | 84-86 | 94-109 | 114-128 | 35.38 | 5.3966 |
| DENV2 | 5-21 | 29-33 | 36-47 | 76-78 | 84-86 | 94-109 | 114-128 | 40 | 5.1234 |
| DENV3 | 5-22 | 29-45 | - | 76-78 | 84-86 | 94-109 | 114-128 | 38.46 | 6.959 |
| DENV4 | 5-21 | 29-45 | - | 76-78 | 84-86 | 94-109 | 114-128 | 38.46 | 7.7112 |
| JEV | 5-22 | 30-45 | - | 77-79 | 85-87 | 96-110 | 115-128 | 49.23 | 6.5367 |
| WNV | 5-22 | 31-48 | - | 77-79 | 85-87 | 96-110 | 116-129 | 47.29 | 7.9211 |
| YFV | 5-21 | 26-45 | - | 76-78 | 84-86 | 95-109 | 113-128 | 30.77 | 6.0231 |
| TBEV | 4-19 | 25-45 | - | 75-77 | 83-85 | 86-108 | 112-128 | 24.39 | 7.2029 |
| MVEV | 5-22 | 30-47 | - | 77-79 | 85-87 | 96-110 | 115-128 | 48.06 | 8.2751 |
| OHFV | 4-18 | 25-46 | - | 75-77 | 83-85 | 86-108 | 111-128 | 24.22 | 7.8924 |
| St Louis EV. | 5-22 | 27-45 | - | 77-79 | 85-87 | 96-110 | 115-128 | 47.69 | 7.0892 |

**Figure 2.**
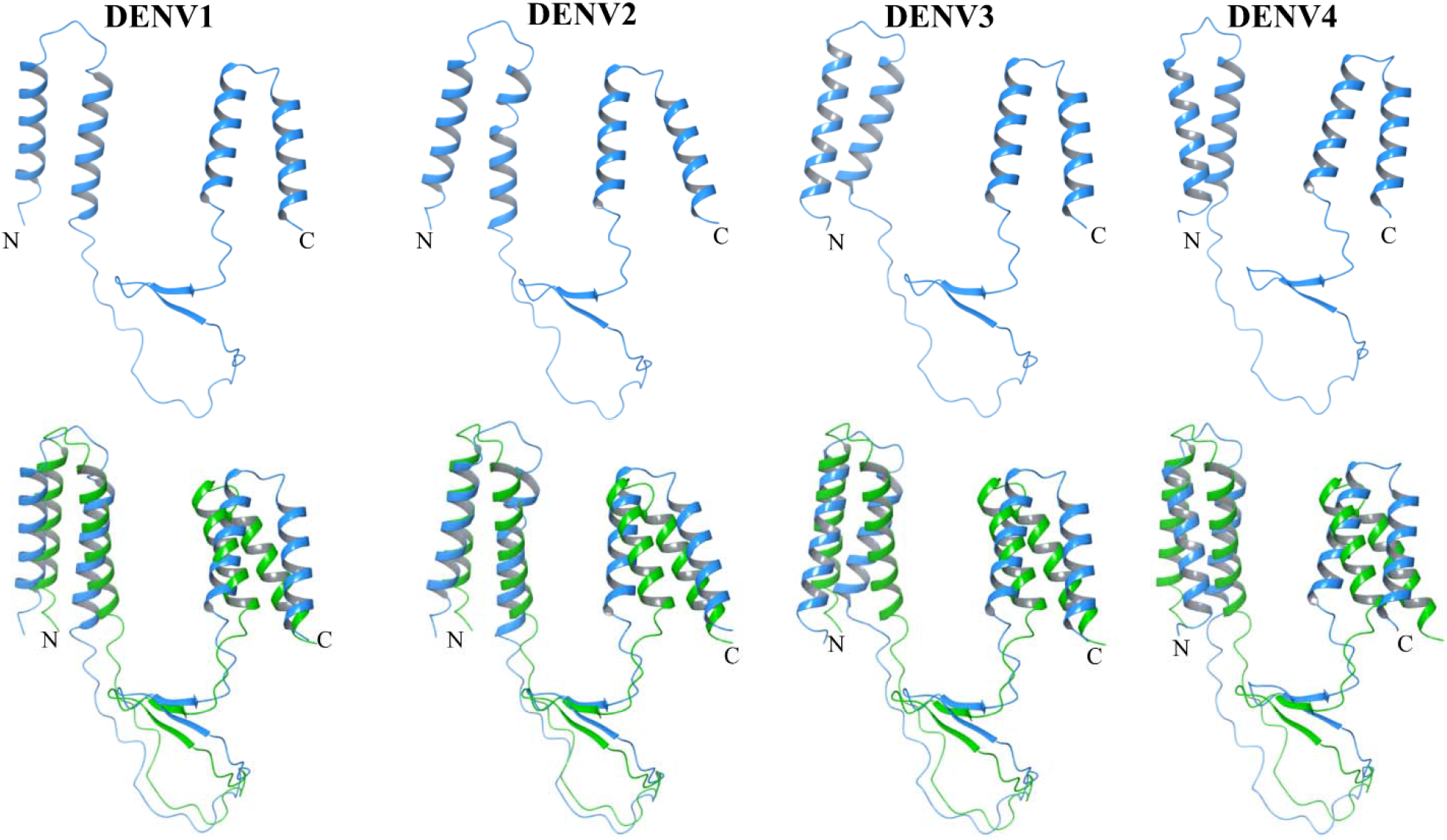
Predicted structure model of NS2B protein of DENV using AlphaFold2 modeling system. The upper panel represents the modeled structure of NS2B protein of different strains of DENV. The lower panel represents the superimposed structure (C-α atoms) of the NS2B protein of the respective DENV strain (blue colored) with the ZIKV NS2B protein (green colored). In all cases, the ranked_0 structure was used to extract the image from the maestro 2018-4 software. N and C indicate the N-terminal and C-terminal ends, respectively.

**Figure 3.**
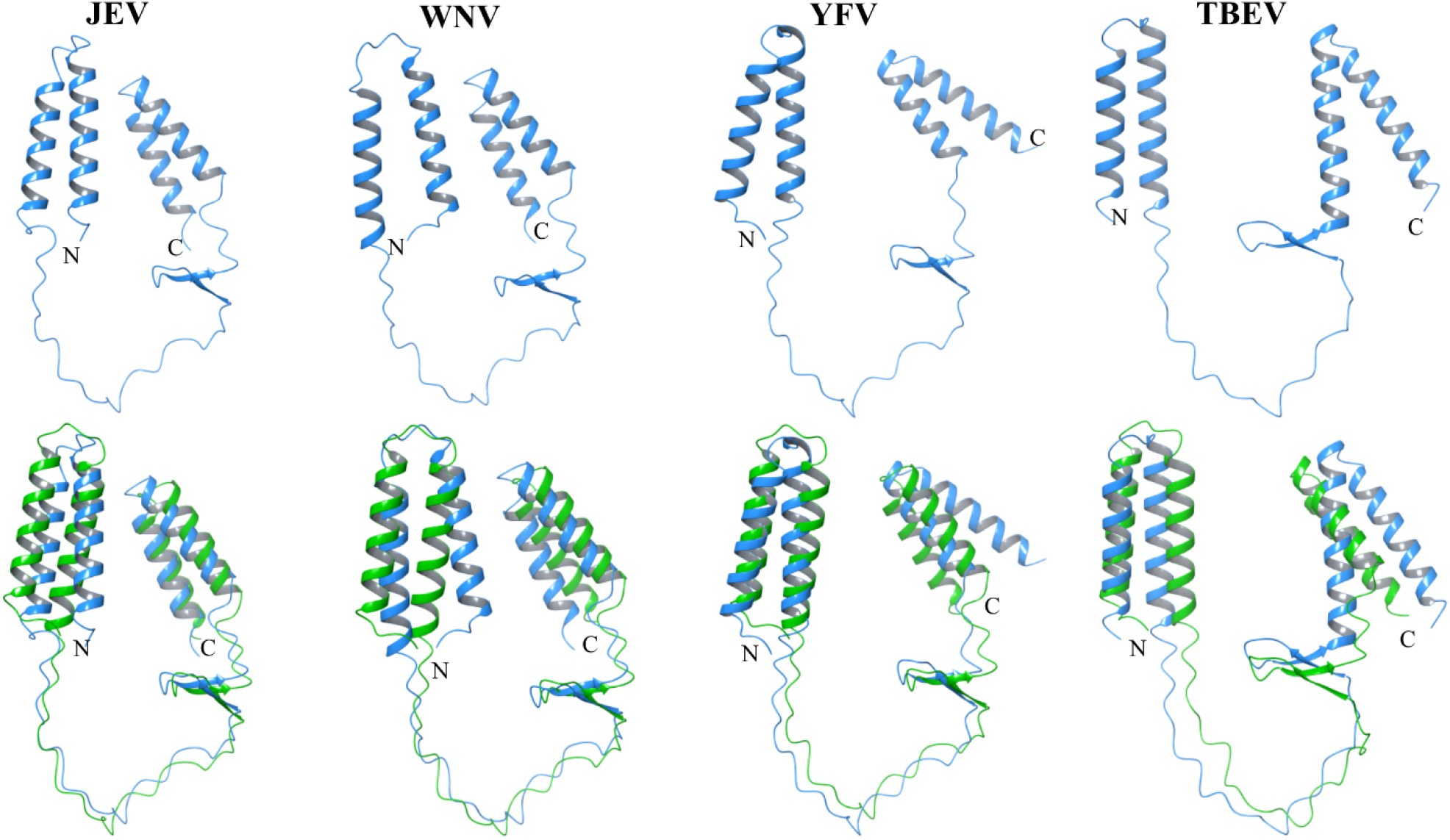
Predicted structure model of NS2B protein of JEV, WNV, YFV, and TBEV using AlphaFold2 modeling system. The upper panel represents the modeled structure of NS2B protein of different strains of respective flaviviruses (JEV, WNV, YFV, and TBEV). The lower panel represents the superimposed (C-α atoms) structure of the NS2B protein of the respective flaviviruses (blue colored) with the ZIKV NS2B protein (green colored). In all cases, the ranked_0 structure was used to extract the image from maestro 2018-4. N and C indicate the N-terminal and C-terminal ends, respectively.

**Figure 4.**
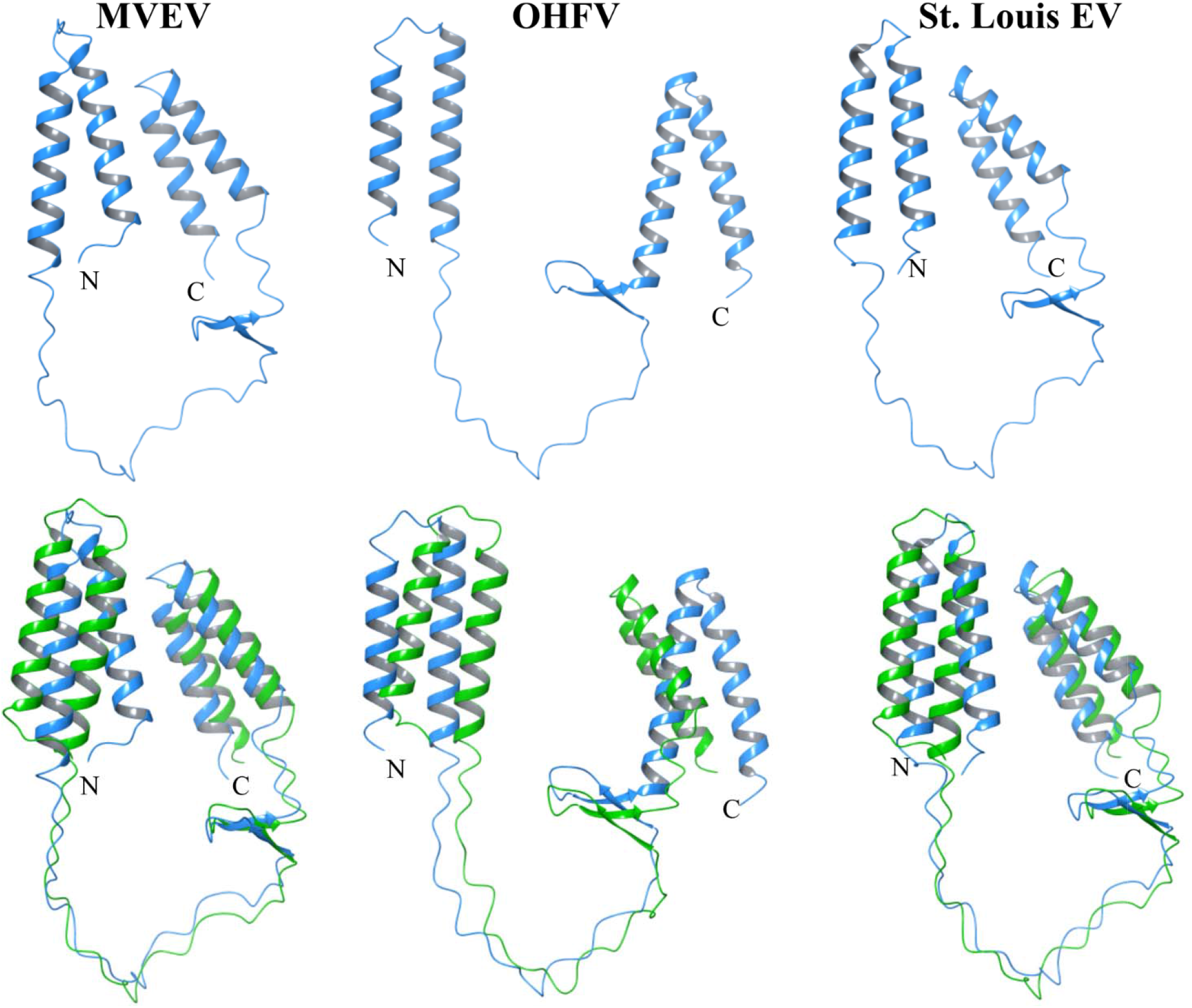
Predicted structure model of NS2B protein of MVEV, OHFV, and St. Louis EV using AlphaFold2 modeling system. The upper panel represents the modeled structure of NS2B protein of different strains of respective flaviviruses (MVEV, OHFV, and St. Louis EV). The lower panel represents the superimposed structure (using C-α atoms) of the NS2B protein of the respective flaviviruses (blue colored) with ZIKV NS2B protein (green colored). In all cases, the ranked_0 structure was used to extract the image from maestro 2018-4. N and C indicate the N-terminal and C-terminal ends, respectively.

The NS2B structure of these flaviviruses contains four helical regions at C-terminal and N-terminal domains and two β-strand in the middle segment (cytosolic domain), visually observed to be similar to the ZIKV NS2B protein structure. However, amid the surprising similarity in predicted structures, a few subtle differences are also present in the modeled structure summarized in Table 1. Notably, the cytosolic domain 45-95 shows a highly disordered structure content, making this segment more flexible.

### 3.2. An insight into the dynamics of ZIKV NS2B full-length protein

Based on the comparative analysis of the NS2B full-length model structure of a few flavivirus members in the previous section, we chose ZIKV NS2B full-length protein to investigate further the secondary structural elements and the dynamics at the residue level.

#### Simulation of NS2B model structure predicted by AlphaFold2

We selected the ranked_0 model structure (Figure 1A) as the template structure and subjected it to MD simulation for 100 ns (Figures 5 & S2). The secondary structure analysis indicates that NS2B has two α helix (residues 5-24 & residues 29-44) at the N-terminal domain (1-44) and two α-helix (residues 94-109 & residues 113-129) at the C-terminal domain (96-130) predominantly stable throughout the simulation. On the other hand, two short β-strands (residues 76-78 & residues 84-86) in the cytosolic domain (45-95) were stable only up to 15 ns and later on disappeared (Figure S2). Besides ordered structures (α-helix and β-strand), NS2B full-length protein also comprises disordered structures (bend, turn, bridge and coil).

**Figure 5.**
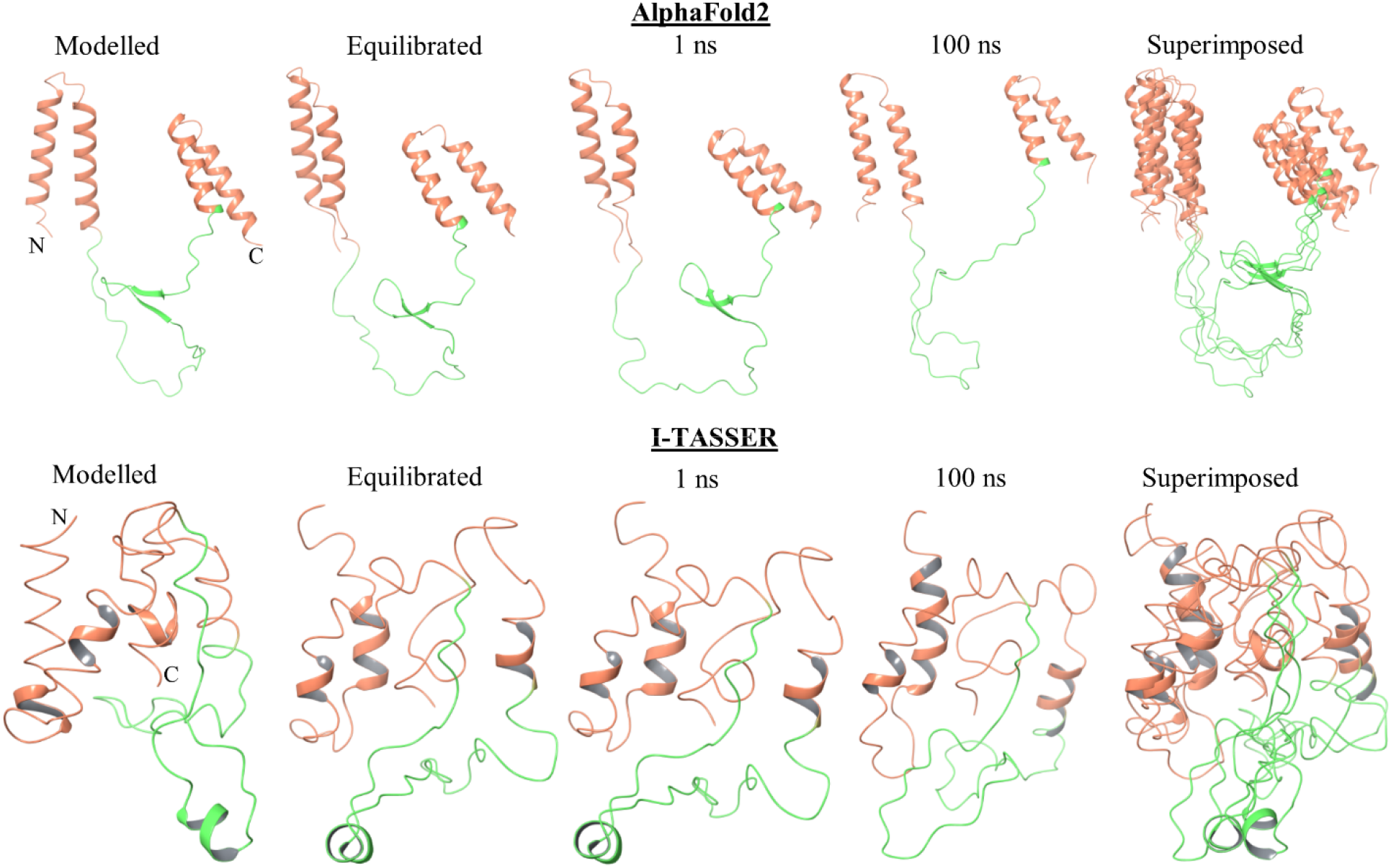
MD simulation of ZIKV NS2B full-length protein. The upper and lower panel represents the snapshot of different time frames trajectory of NS2B full-length protein (structure predicted by both I-TASSER and AlphaFold2 modeling system) during 100 ns MD simulation. The modeled structure was placed inside a bilayer membrane consisting of 150 molecules of DOPC lipids. The green-colored segment represents the cytosolic domain (residues 49-95) of NS2B.

#### Simulation of NS2B model structure predicted by I-TASSER

We selected the highest scoring (C-score -3.71; see Figure 1B) model structure as template structure and subjected it to MD simulation for 100 ns (Figures 5 & S2). The secondary structure analysis indicates that NS2B has two α helix (residues 1-20 & residues 30-40) at the N-terminal domain (1-44) and one α-helix (residues 115-125) at the C-terminal domain (96-130) that is predominantly stable throughout the simulation. On the other hand, one α-helix (residues 63-79) appeared in the cytosolic domain (45-95), stable only up to 30 ns, and later on, disappeared (Figure S2).

Very few residues contributed to the β-strand that appeared in very few time frames distributed non-uniformly. Besides ordered structures (α-helix and β-strand), NS2B full-length protein also comprises disordered structures (bend, turn, bridge and coil). The residues of the cytosolic domain (49-95) show higher RMSF than the N-terminal and C-terminal domain (Figure S2) in both model structures. Higher fluctuating residues in the cytosolic domain result from the disordered structure being more flexible and dynamic. Overall simulation results suggest that the cytosolic domain of NS2B full-length protein has a mostly disordered structure in line with the previous report [10].

### 3.3. NS2B cytosolic domain (residues 49-95) shows extensive flexibility

The cytosolic domain of NS2B protein has a significant contribution as a co-factor for its partner protein [2,7,14]. Therefore, we investigate the dynamics of this domain in isolation to see how many different conformations it can attain under changing environmental conditions. At first, we modeled the NS2B cytosolic domain (49-95) using I-TASSER modeling system (Figure 6A). Then, to further support the I-TASSER result, we also modeled NS2B (49-95) structure using the AlphaFold2 modeling system (Figure 6B).

**Figure 6.**
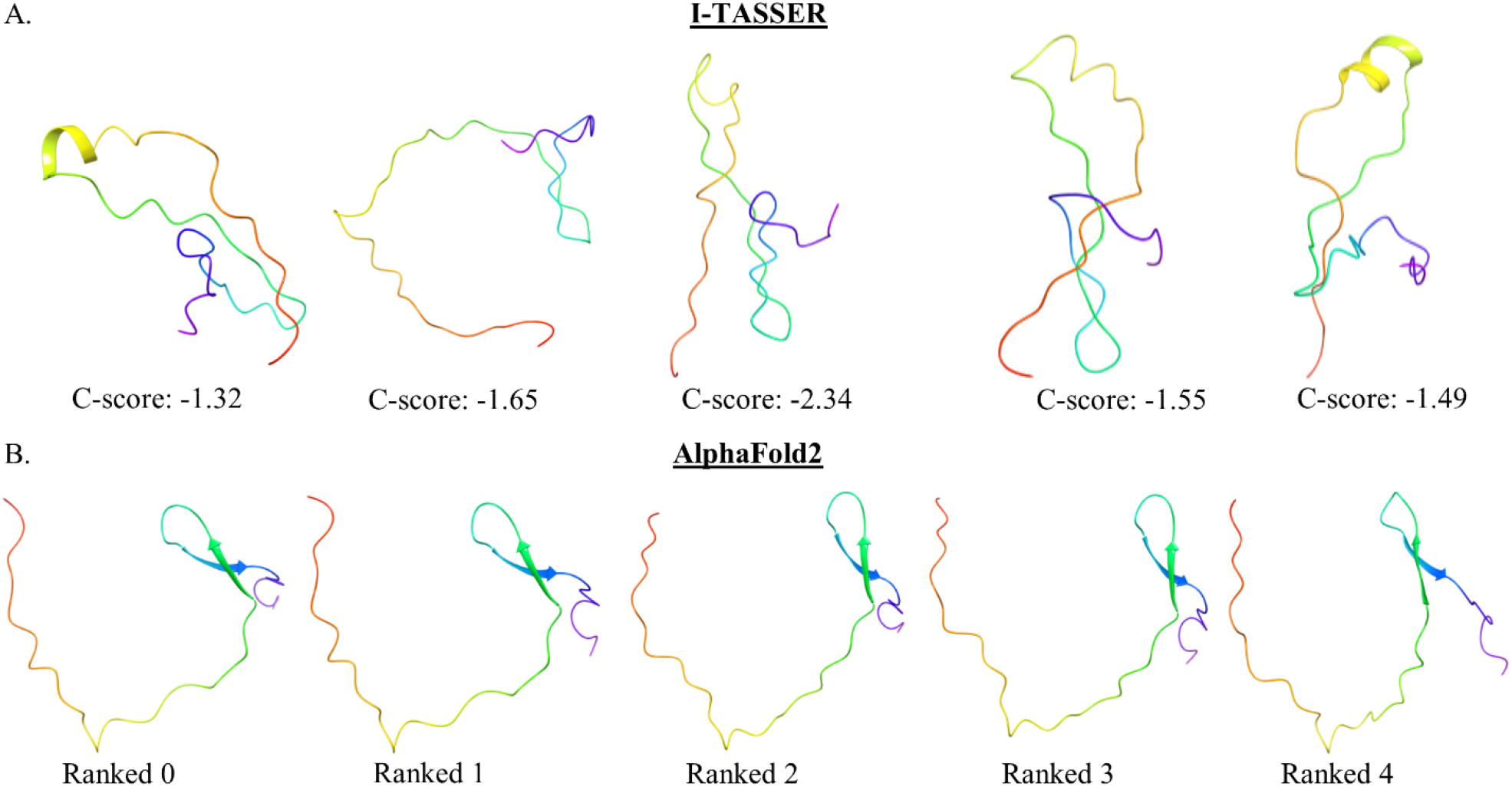
Structure modeling of ZIKV NS2B cytosolic domain (residues 49-95). (A) Model structure of ZIKV NS2B (49-95) from I-TASSER. The structure with a C-score of -1.32 is used as a template and further subjected to MD simulation. The red-colored terminal is the N-terminal (residue 49^th^), and the purple-colored terminal is the C-terminal (residue 95^th^) of the NS2B (49-95). (B) Model structure of ZIKV NS2B (49-95) from AlphaFold2. The ranked_0 structure is used as a template to analyze the structural elements. The red-colored terminal end is the N-terminal (residue 49^th^), and the purple-colored terminal end is the C-terminal (residue 95^th^) of the NS2B (49-95).

The structure with a C-score of -1.32 in the I-TASSER system and the ranked_0 structure in the AlfaFold2 system was used as template structure. The NS2B (49-95) contains one α-helix (18-21) predicted by I-TASSER system. On the other hand, the AlphaFold2 model suggests two β-strand (28-30 & 36-38). Therefore, NS2B (49-95) modeled structure has a high content of disordered structure, which is in line with the previously determined secondary structure of NS2B (49-95) as a disordered type protein in isolation using a disordered prediction server and the CD spectroscopy [11]. Furthermore, the selected model structures of NS2B (49-95) from both predicting systems were subjected to MD simulation (Figure S3) for 500 ns in water. Further, the last frame at 500 ns was used as a template structure to run simulation for a longer time (1000 ns).

In figure 7, I-TASSER modeled NS2B (49-95) structure in water showed predominantly two β-strands at the c-terminal (residue 72-78 and residues 84-90) of this peptide which is contributed by 14 residues only (Figure 7A). The remaining 33 residues contributed to the disordered structure towards the N-terminal of this peptide. This peptide also forms helices for a short duration between 0 to 50 ns and 200 to 275 ns at residues 63 to 68 and 65 to 70, respectively (Figure 7B). Moreover, it shows minimum fluctuating RMSD and RMSF (Figure 7B). Similarly, AlphaFold modeled NS2B (49-95) structure in water also showed predominantly β-strands in the peptide (Figure 7C & D). Moreover, it shows minimum fluctuating RMSD and RMSF (Figure 7D) correlate well with the simulation of I-TASSER modeled structure. The simulation analysis indicates that the NS2B cytosolic domain (49-95) is disordered in isolation as a peptide segment and shows a good agreement with the previously characterized NS2B cytosolic domain (49-95) as a disordered peptide [11].

**Figure 7.**
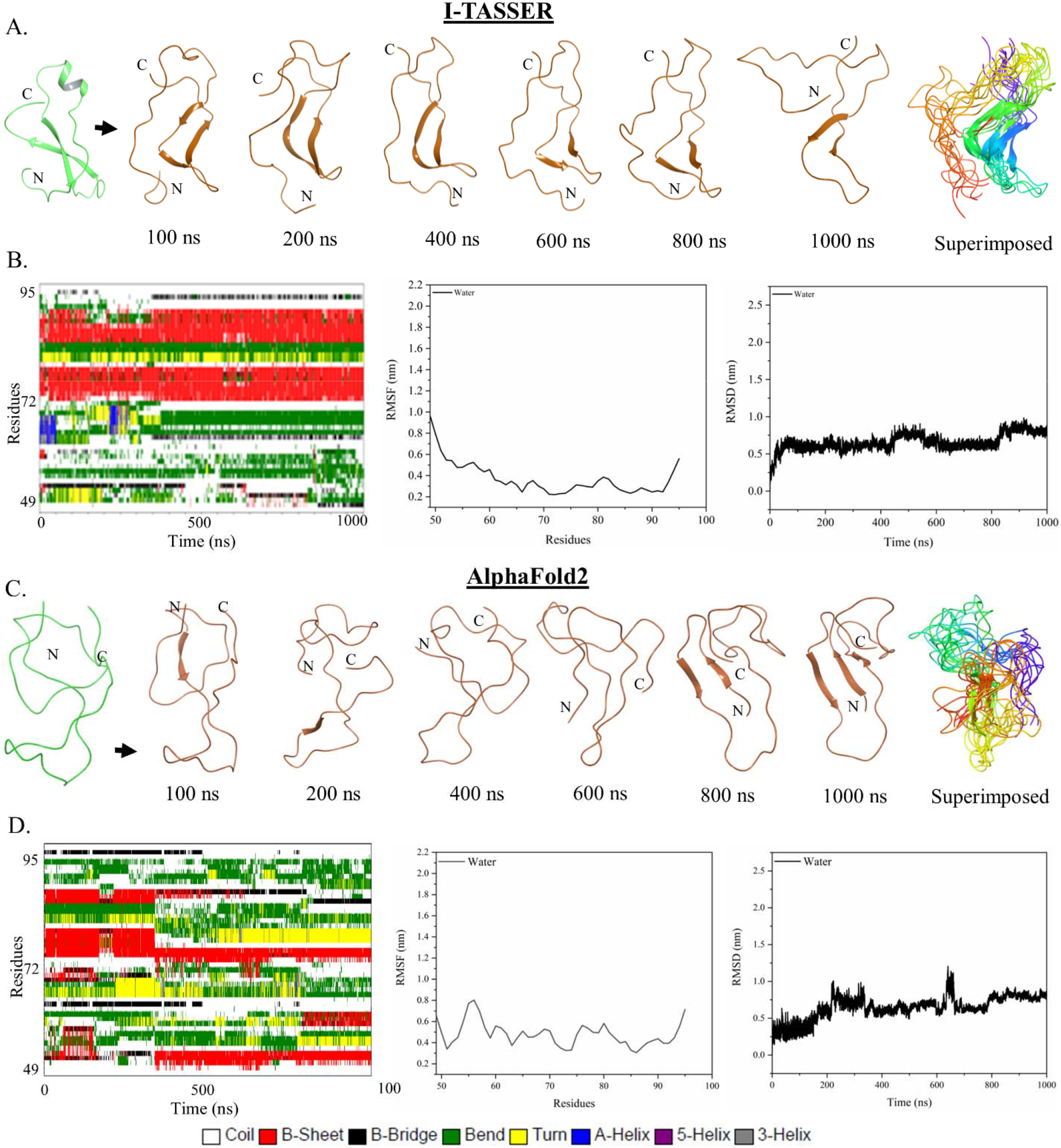
MD simulation of NS2B cytosolic domain (residues 49-95) in water. (A) & (C) represents a snapshot of the different time frames trajectory of NS2B (residues 49-95) modeled structure, predicted by I-TASSER and AlphaFold2, respectively. The red-colored end is the N-terminal (49^th^ residue), and the purple-colored end is the C-terminal (95^th^ residue) in the superimposed structures. (B) & (D) represents the MD simulation parameter of NS2B (residue 49-95) modeled structure by I-TASSER and AlphaFold2, respectively. 1^st^ panel represents the contribution of NS2B (residues 49-95) residues to the secondary structure where red indicate the β-strand at the respective region of this peptide, and blue indicates the helices at th corresponding region of the peptide. 2^nd^ and 3^rd^ panels show the RMSF of each residue and RMSD during the simulation period.

### 3.4. TFE induces conformational changes in NS2B cytosolic domain (residues 49-95)

When the NS2B (49-95) peptide (modeled by I-TASSER) was subjected to simulation in TFE, two β-strands were present at the same position at the C-terminal of the peptide, as observed in the case of water. TFE also induces α-helices at the N-terminal of the peptide (residues 55-60 and residues 63-70) without affecting two β-strand at the C-terminal (Figure 8A). Overall, the simulation analysis indicates that NS2B (49-95) in TFE possesses higher α-helical content than water. The change in conformation to this peptide in the presence of TFE shows high RMSD or RMSF compared to the structure in water (Figure 8B). This simulation data suggests that TFE induces conformational changes at the N-terminal region of the peptide without affecting the β-strand at C-terminal. NS2B (49-95) peptide (modeled by AlphaFold2) in TFE shows four β-strands (residues 52-54, residues 68-70, residues 73-79, and residues 82-86; Figure 8C & D). TFE also induces short α-helices towards the C-terminal of the peptide (residues 67-70 and residues 75-79) for very few nanoseconds (Figure 8D). The structure shows high RMSD or RMSF compared to the structure in water (Figure 8D). The simulation analysis indicates that NS2B (49-95) in TFE induces disorder to ordered structural transition.

**Figure 8.**
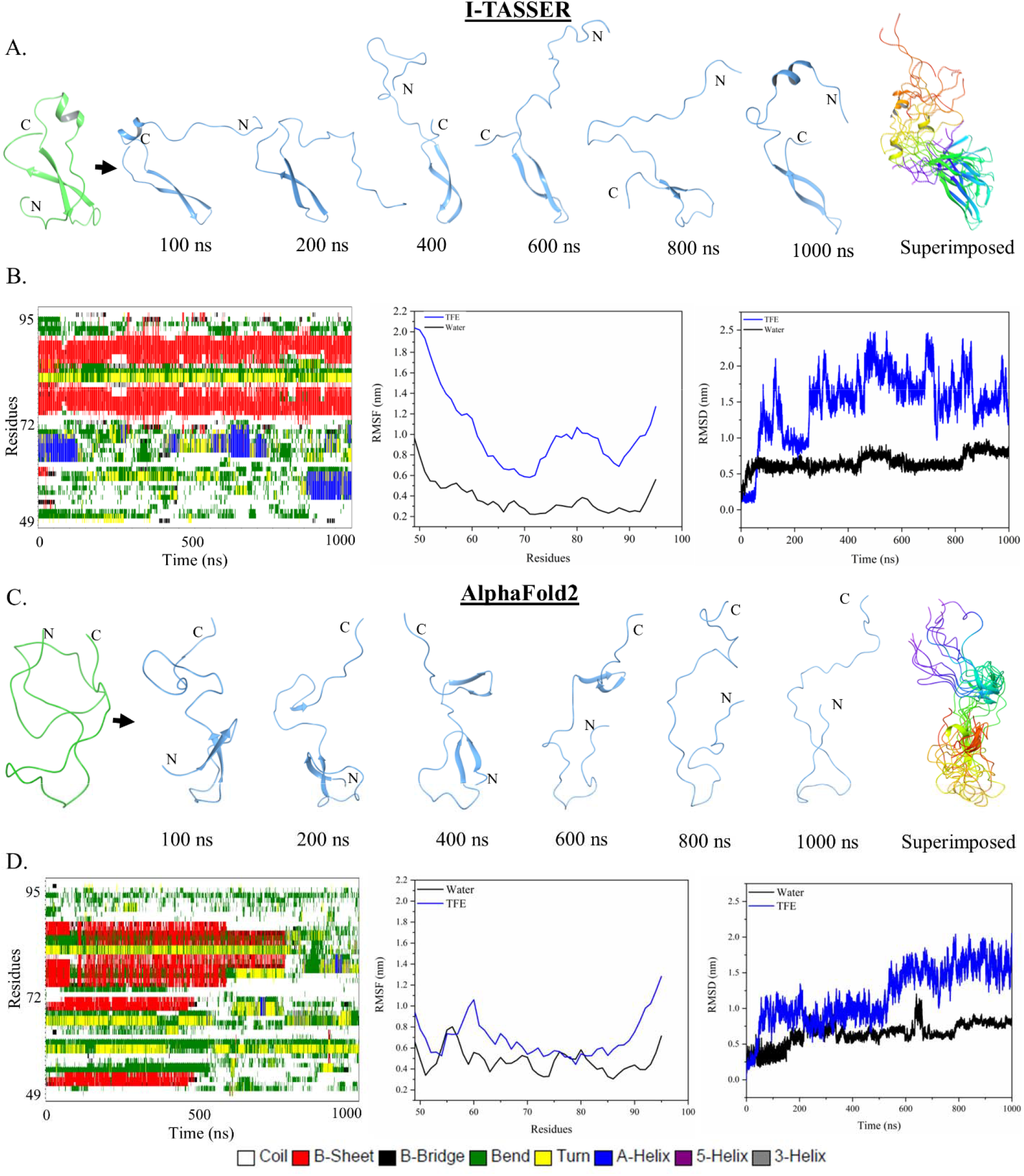
MD simulation of NS2B cytosolic domain (residues 49-95) in TFE. (A) & (C) represents a snapshot of the different time frames trajectory of NS2B (residues 49-95) modeled structure, predicted by I-TASSER and AlphaFold2, respectively. The red-colored end is the N-terminal (49^th^ residue), and the purple-colored end is the C-terminal (95^th^ residue) in the superimposed structures. (B) & (D) represents the MD simulation parameter of NS2B (residues 49-95) modeled structure by I-TASSER and AlphaFold2, respectively. 1^st^ panel represents the contribution of NS2B (residues 49-95) residues to the secondary structure where red indicates the β-strand at the respective region of this peptide, and blue indicates the helices at the corresponding region of the peptide. 2^nd^ and 3^rd^ panels show the RMSF of each residue and RMSD during the simulation period.

Further, the finding of the simulation study correlates well with the finding of the CD spectroscopy (Figure 9). The CD spectrum of the peptide NS2B (49-95) in an aqueous solution was found to have a signature minimum at 198 nm, which showed characteristic features of random coil/disordered peptide/protein. But in the presence of TFE, NS2B (49-95) peptide shows a remarkable change in the shape of the CD spectra (Figure 9A). The CD spectra at 60-70% TFE show a signature minimum near 208 nm and 222 nm, a typical protein characteristic with a high content of α-helices. Further, the change in conformation and transition state of th peptide was evaluated by qualitative analysis of ellipticity at 198 nm vs. 222 nm, showing a gain of an alpha helix in the presence of TFE (Figure 9B).

**Figure 9.**
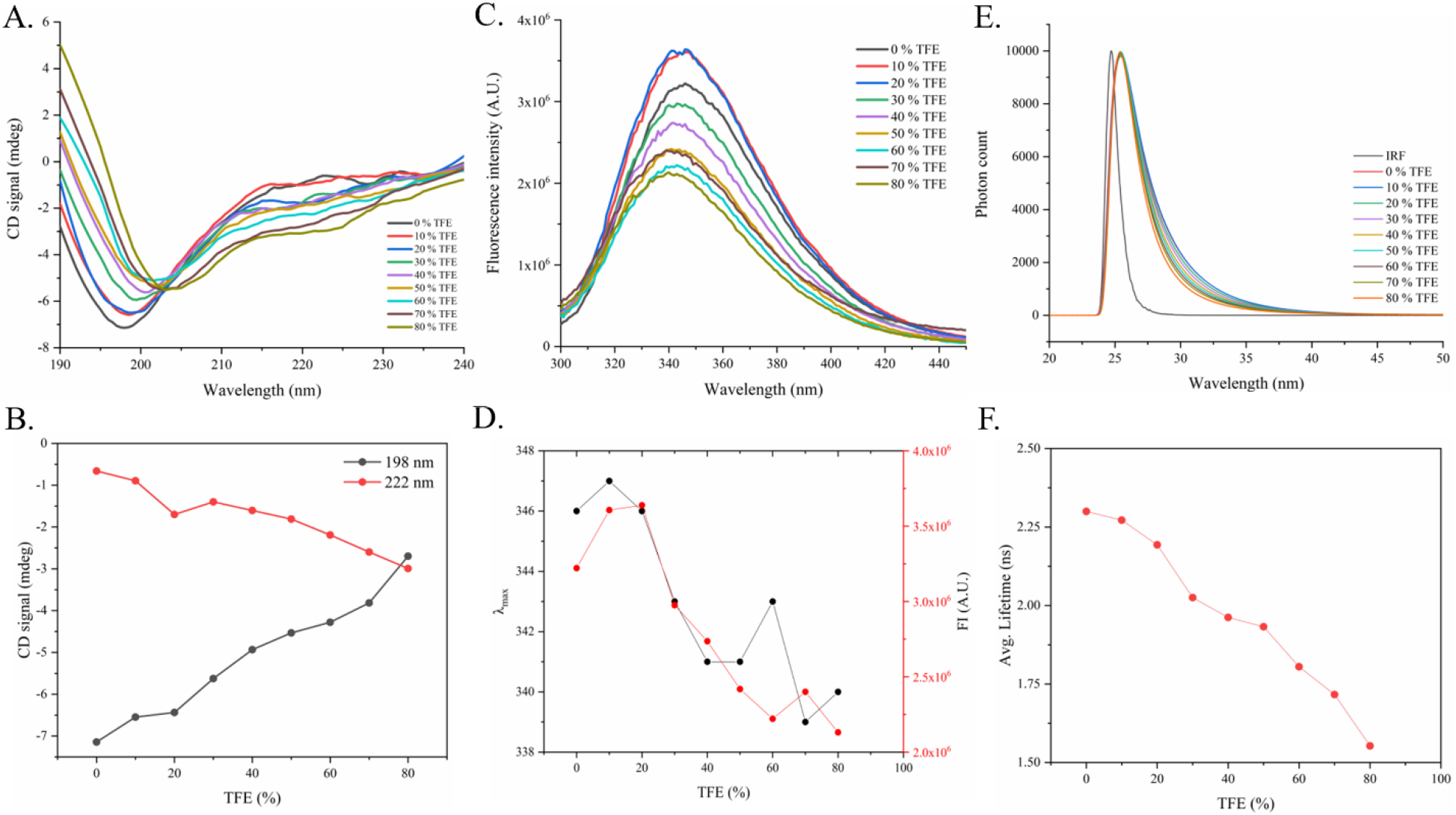
TFE induces conformational changes in NS2B cytosolic domain (residues 49-95). (A) Far UV-CD spectra of the NS2B peptide (49-95) in the presence of TFE, respectively, at 25°C. (B) A plot of the CD signal at 198 nm and 222 nm as the function of TFE. (C) The fluorescence spectra of NS2B (49-95) peptide in the presence of TFE at 20°C. (D) Emission maxima of NS2B (49-95) peptide as the function of TFE. (E) Fluorescence lifetime spectra of NS2B (49-95) peptide in TFE. (F) Average lifetime of NS2B (49-95) peptide as the function of TFE.

Further, we have investigated the effect of TFE on the tertiary structure of the NS2B (49-95) peptide. Trp was used as an intrinsic probe to assess any change in tertiary structure conformation influenced by the surrounding environment. The Trp fluorescence studies of the NS2B peptide show an emission maximum at 347 nm in the presence of phosphate buffer. In the presence of TFE, the intensity at emission maxima shows a decreasing trend as the function of TFE (Figure 9C) with a blue shift (Figure 9D) due to an increased non-polarity of TFE compared to water. At the same time, the blue shift and the fluorescence intensity change may be due to different orientations of the Trp indole ring, which may show a different mode of interacting behavior in the presence of either TFE or water. Tertiary structure conformation dynamics were further investigated by fluorescence lifetime measurement. The decay data were fitted in a three-term exponential model with best-fit values of χ2 near one, as shown in Figure 9E. In an aqueous solution, the average lifetime of NS2B (49-95) peptide was observed to be 2.30 ns, whereas, in TFE (80%), the average lifetime was 1.55 ns (Table 2). The average lifetime shows a decreasing trend with the function of TFE (Figure 9F) correlates well with the fluorescence emission spectra result. The average lifetime and fluorescence emission spectra suggest the TFE induces compaction of NS2B (49-95) overall structure, strengthening the CD spectroscopy results where TFE induces α-helix conformation.

**Table 2.** Lifetime parameters associated with the NS2B (residues 49-95) in the presence of TFE and SDS. τ is the average lifetime.

| Component | Conc. | $\tau$ (ns) | $\chi^2$ |
| --- | --- | --- | --- |
| NS2B (49-95) | 5 μM | 2.30 | 1.24 |
| TFE | 10 % | 2.27 | 1.34 |
|  | 20 % | 2.19 | 1.14 |
|  | 30 % | 2.03 | 1.23 |
|  | 40 % | 1.96 | 1.21 |
|  | 50 % | 1.93 | 1.24 |
|  | 60 % | 1.80 | 1.21 |
|  | 70 % | 1.72 | 1.27 |
|  | 80 % | 1.55 | 1.26 |
| SDS | 1 mM | 2.12 | 1.19 |
|  | 5 mM | 1.93 | 1.27 |
|  | 25 mM | 1.74 | 1.26 |
|  | 100 mM | 1.72 | 1.24 |

### 3.5. Effect of SDS, molecular crowding, and temperature on the structure of NS2B cytosolic domain (residues 49-95)

Further, we have investigated the conformational dynamics of NS2B cytosolic domain (49-95) peptides in SDS and crowded environments. The macromolecule concentration inside the cell is very high, thus has excluded volume effect and dehydration, which significantly influence protein structures [23,24]. Previously Ficoll and PEG were used as molecular crowding agents [25].

The change in the secondary structure conformation in NS2B cytosolic domain (49-95) was determined by Far UV-CD spectra, where both SDS and molecular crowding agent (PEG8000 & Ficoll 70) do not exhibit any effect on the shape of the spectra (Figures 10A, 11A and 11B). Furthermore, analysis of fluorescence emission spectra shows a change in the intensity at the emission maxima in SDS and molecular crowder. For example, at 5 mM and 25 mM SDS, the fluorescence intensity is higher than its negative control (i.e., absence of SDS). On the other hand, at a higher concentration of 100 mM, the intensity is almost equal, but the emission maxima show a blue shift (Figure 10B). Furthermore, the lifetime measurement (Figure 10C) revealed that the average lifetime of NS2B cytosolic domain (49-95) decreases with an increase in SDS concentration (Figure 10D).

**Figure 10.**
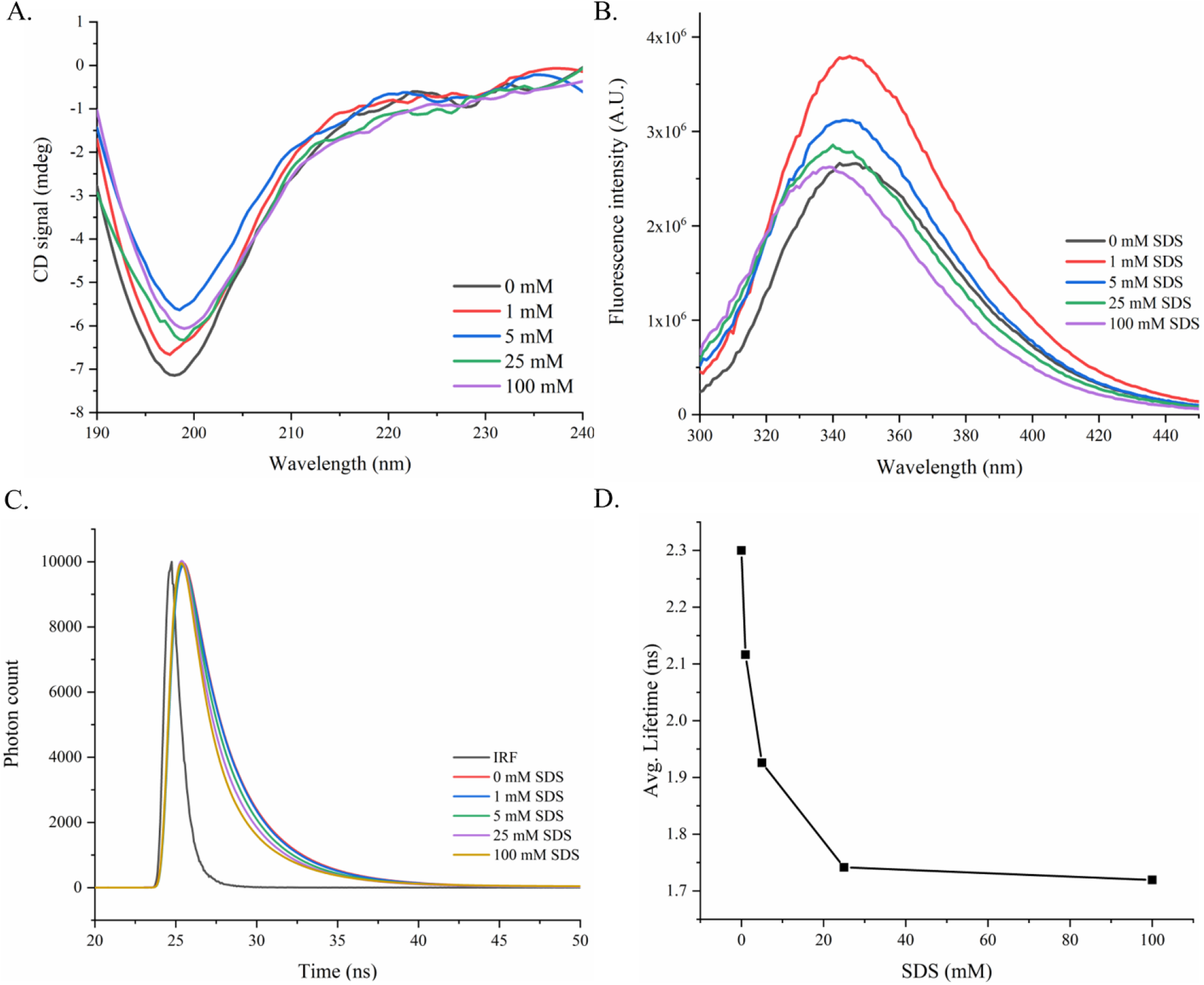
Effect of SDS on the structure of NS2B cytosolic domain (residues 49-95). (A) Far UV-CD spectra of NS2B cytosolic domain (49-95) in SDS presence show disordered type peptide spectra. (B) Fluorescence spectra of NS2B cytosolic domain (49-95) in the presence of an increasing concentration of SDS. (C) Fluorescence lifetime measurement of NS2B cytosolic domain (49-95) in the presence of SDS. (D) Average lifetime of NS2B cytosolic domain (49-95) as the function of SDS.

**Figure 11.**
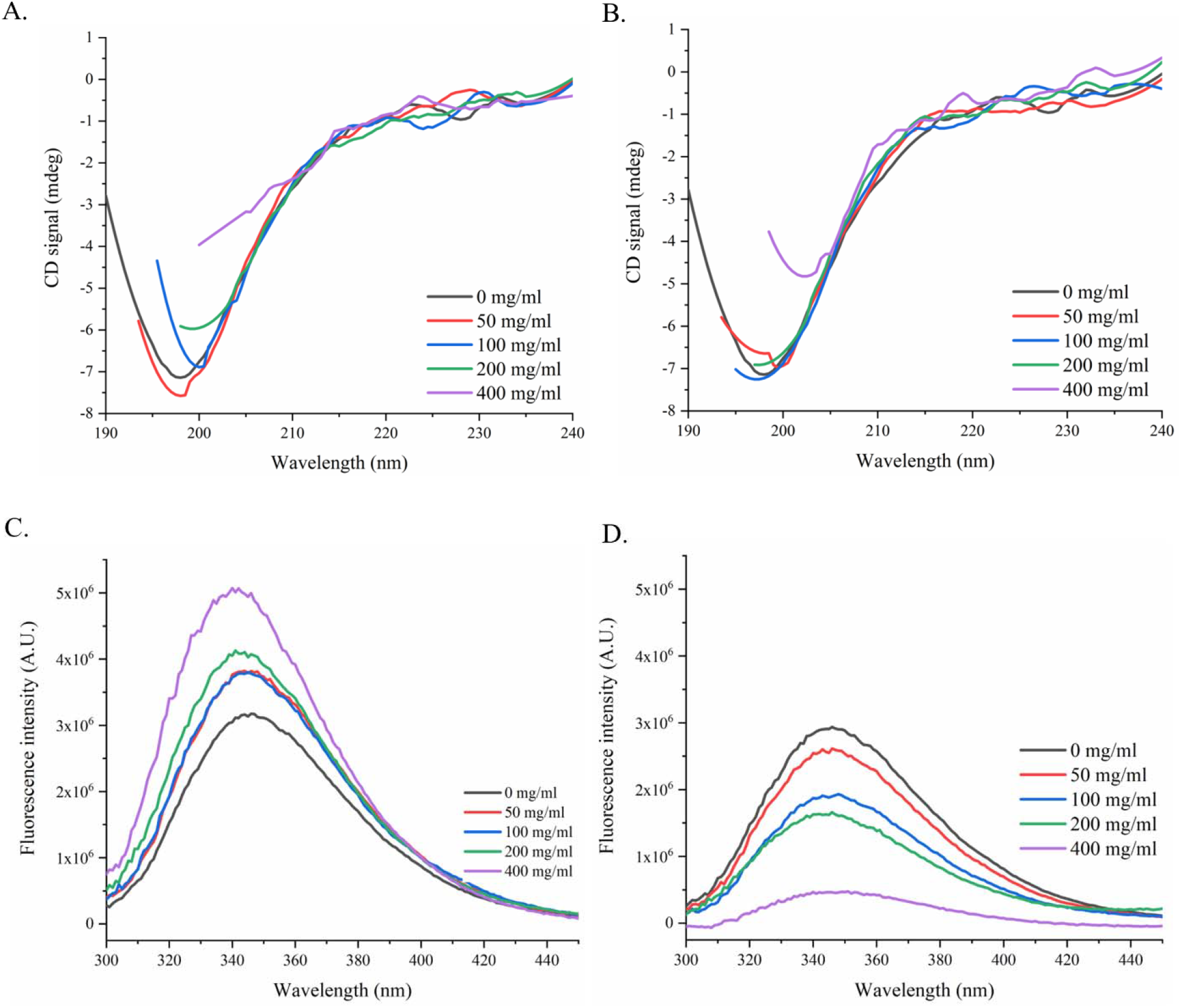
Effect of PEG and Ficoll on the structure of NS2B cytosolic domain (residues 49-95). (A) and (B) represents Far UV-CD spectra of NS2B cytosolic domain (49-95) in the presence of PEG8000 and Ficoll70, respectively, showing the disordered protein spectrum. (C) and (D) represents fluorescence emission spectra of NS2B cytosolic domain (49-95) in th presence of PEG8000 and Ficoll70, respectively.

The emission fluorescence intensity increases as a function of PEG8000 and decreases as a function of Ficoll70. The change in the fluorescence intensity maxima indicates that the presence of these agents affects the dynamics of the tertiary structure of NS2B cytosolic domain (49-95) (Figures 11C & 11D). Further, we monitored the Far UV-CD spectra (Figure 12) of NS2B cytosolic domain (49-95) with varying temperatures (10°C-95°C). The change in the shape and signature minima of the NS2B cytosolic domain (49-95) spectra as the function of temperature (observed a partial contraction at a higher temperature) is very similar to the spectra of the disordered type peptide reported previously [16,26].

**Figure 12.**
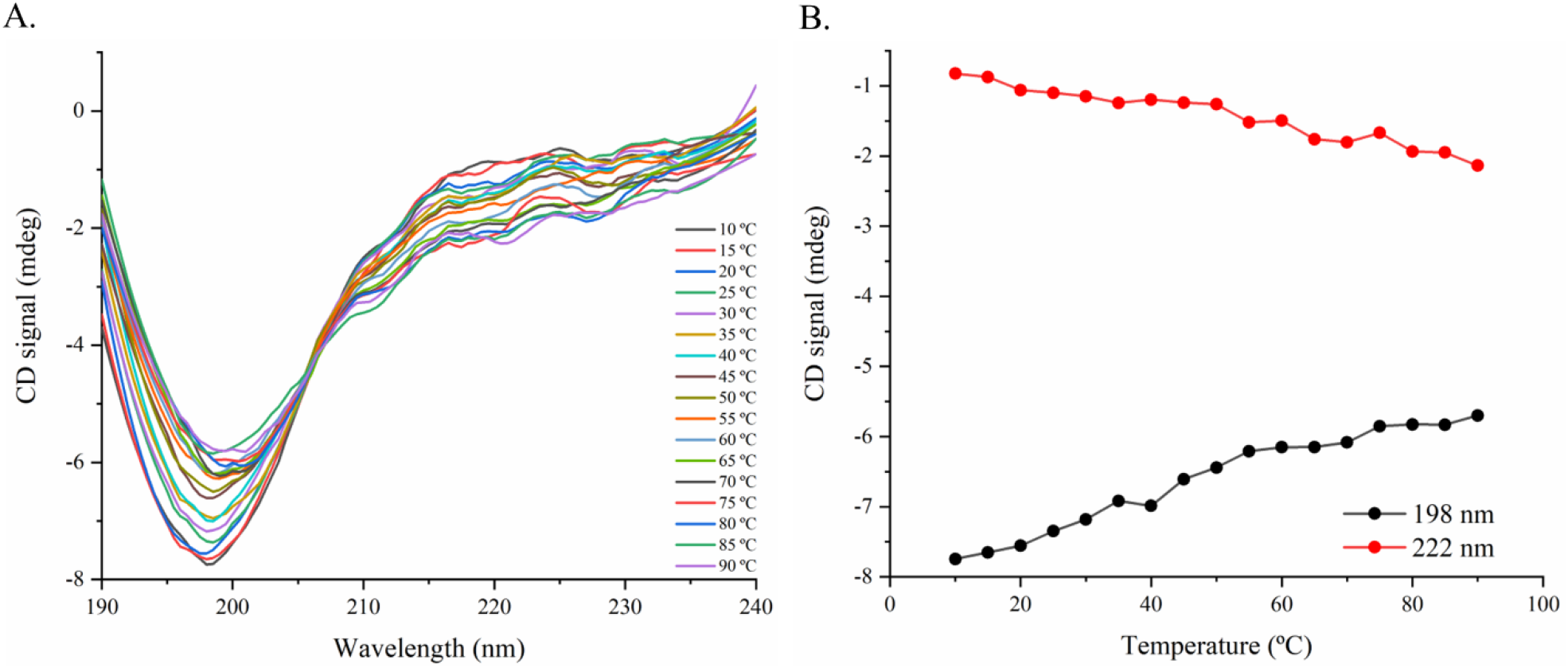
Effect of temperature on the NS2B cytosolic domain (residues 49-95). (A) Far UV-CD spectra of NS2B cytosolic domain (49-95) at varying temperatures (10°C to 9°C) with 5°C interval showing typical disorder-type protein spectra. (B) CD signal at 198 and 222 nm as the function of temperature.

## 4. Discussion

NS2B acts as a co-factor for the activity of the NS3 protease in the flaviviruses [27]. Together, these proteins form NS2B-NS3 protease complex needed for cleaving a single polyprotein encoded by flaviviruses [27,28], where only the NS2B cytosolic domain is sufficient for the functional activity of NS3 protease. On the other hand, the N-terminal and C-terminal transmembrane domains provide membrane anchors to the NS3 protease [2,6,9]. In recent decades, many X-ray crystal structures of NS2B-NS3 protease with high resolution have been determined in complex with inhibitors molecules where only the NS2B cytosolic domain has been characterized in detail [7,8]. Still, the information about the dynamics and the structural elements of NS2B full-length protein is minimal [2,6,10]. So, in this article, we have first analyzed the alphafold2 modeled structure of ZIKV NS2B and compared it with the NS2B structure of other members of flaviviruses, viz. DENV, JEV, WNV, YFV, OHFV, St. Louis EV, MBEV and TBEV. Despite only 25-48% sequence identity with the ZIKV NS2B full-length protein, all the flavivirus members show surprising similarities fold-wise with subtle differences in the modeled structures. The NS2B full-length structure suggests four helical regions at the N-terminal and C-terminal transmembrane domains and two β-strands at the middle segment (cytosolic domain). The middle segment in these viruses has a mostly unstructured (disordered) region that makes this region highly flexible. As a result, it may attain multiple conformations to interact with partner proteins and other functions. Furthermore, in ZIKV, the NS2B cytosolic domain wraps around the NS3 protease and forms β-strands near the protease’s active site [1], forming the NS2B-NS3 protease complex. The cytosolic domain (residues 49-95) in this complex forms four β-strand [7]. Similarly, in the dengue virus (share 55-57 % amino acid sequence identity with ZIKV)[29], the NS2B cytosolic domain (48-100) in the protease complex (PDB ID: 3U1I) [30] forms four β-strands with no helix, but the isolated NS2B cytosolic domain (48-100) composed more of helix [31]. Therefore, the evidence suggests that upon interaction with the NS3 protease, the NS2B cytosolic domain adopts more β-strands. On the contrary, the ZIKV NS2B cytosolic domain (49-95) in isolation was previously identified as an intrinsically disordered (high content of unstructured region as revealed by CD spectra) peptide [11]. The multiple conformations of the Flavivirus NS2B cytosolic domain may correlate well with previously reported literature that the organic solvent induces a conformational change (α-helix and β-strand) in various disordered proteins [26,32,33]. In the ZIKV NS2B cytosolic domain (49-95), the TFE induces conformational transition at the N-terminal region without affecting the two β-strand at the C-terminal region. It may suggest that the C-terminal region is very flexible and dynamic, requiring interaction with the NS3 protease to adopt an additional two β-strand. We also believe that apart from the known co-factor role of the NS2B cytosolic region, it may use its alpha-helical conformation for some unknown function.

## Authors contribution

RG designed the study. AK performed the majority of experiments, wrote the first draft, and analyzed the data with input from RG. RG edited the manuscript and acquired the funding. AK and PK performed the computational part. PMM helped in CD experiment for TFE. All authors approved the final draft.

## Acknowledgments

We want to thank IIT Mandi for providing research space and facilities. This work is supported by the Indian Council of Medical Research (52/04/2020/BIO/BMS) grant to RG. RG is also grateful Science and Engineering Research Board (SERB) grant, India (CRG/2019/005603); IYBA Award (BT/11/IYBA/2018/06) grant, Department of Biotechnology (DBT), India; Indian Council of Medical Research (58/6/2020/PHA/BMS), and MHRD-SPARC (SPARC/2018-2019/P37/SL).

## References

[1] R. Hilgenfeld, J. Lei, L. Zhang, The Structure of the Zika Virus Protease, NS2B/NS3<Superscript>pro</Superscript>, Adv. Exp. Med. Biol. 1062 (2018) 131–145. doi:10.1007/978-981-10-8727-1_10.

[2] H. Xing, S. Xu, F. Jia, Y. Yang, C. Xu, C. Qin, L. Shi, Zika NS2B is a crucial factor recruiting NS3 to the ER and activating its protease activity, Virus Res. 275 (2020) 197793. doi:10.1016/J.VIRUSRES.2019.197793.

[3] Y. Li, Q. Li, Y.L. Wong, L.S.Y. Liew, C. Kang, Membrane topology of NS2B of dengue virus revealed by NMR spectroscopy, Biochim. Biophys. Acta - Biomembr. 1848 (2015) 2244–2252. doi:10.1016/J.BBAMEM.2015.06.010.

[4] M. León-Juárez, M. Martínez-Castillo, G. Shrivastava, J. García-Cordero, N. Villegas-Sepulveda, M. Mondragón-Castelán, R. Mondragón-Flores, L. Cedillo-Barrón, Recombinant Dengue virus protein NS2B alters membrane permeability in different membrane models, Virol. J. 13 (2016) 1–11. doi:10.1186/S12985-015-0456-4/FIGURES/7.

[5] X.-D. Li, C.-L. Deng, H.-Q. Ye, H.-L. Zhang, Q.-Y. Zhang, D.-D. Chen, P.-T. Zhang, P.-Y. Shi, Z.-M. Yuan, B. Zhang, Transmembrane Domains of NS2B Contribute to both Viral RNA Replication and Particle Formation in Japanese Encephalitis Virus, J. Virol. 90 (2016) 5735–5749. doi:10.1128/JVI.00340-16/ASSET/EAC56B7E-8CC3-4F3D-9259-D3434B7EF765/ASSETS/GRAPHIC/ZJV9991817070010.JPEG.

[6] D. Aguilera-Pesantes, M.A. Méndez, Structure and sequence based functional annotation of Zika virus NS2b protein: Computational insights, Biochem. Biophys. Res. Commun. 492 (2017) 659–667. doi:10.1016/J.BBRC.2017.02.035.

[7] J. Lei, G. Hansen, C. Nitsche, C.D. Klein, L. Zhang, R. Hilgenfeld, Crystal structure of zika virus ns2b-ns3 protease in complex with a boronate inhibitor, Science (80-.). 353 (2016) 503–505. doi:10.1126/SCIENCE.AAG2419/SUPPL_FILE/LEI-SM.PDF.

[8] Z. Zhang, Y. Li, Y.R. Loh, W.W. Phoo, A.W. Hung, C.B. Kang, D. Luo, crystal structure of unlinked NS2B-NS3 protease from Zika virus, Science (80-.). 354 (2016) 1597–1600. doi:10.1126/SCIENCE.AAI9309/SUPPL_FILE/AAI9309-ZHANG-SM.PDF.

[9] S. Clum, K.E. Ebner, R. Padmanabhan, Cotranslational Membrane Insertion of the Serine Proteinase Precursor NS2B-NS3(Pro) of Dengue Virus Type 2 Is Required for Efficient in Vitro Processing and Is Mediated through the Hydrophobic Regions of NS2B *, J. Biol. Chem. 272 (1997) 30715–30723. doi:10.1074/JBC.272.49.30715.

[10] E.Y. Ng, Y.R. Loh, Y. Li, Q. Li, C.B. Kang, Expression, purification of Zika virus membrane protein-NS2B in detergent micelles for NMR studies, Protein Expr. Purif. 154 (2019) 1–6. doi:10.1016/J.PEP.2018.09.013.

[11] P.M. Mishra, V.N. Uversky, R. Giri, Molecular Recognition Features in Zika Virus Proteome, J. Mol. Biol. 430 (2018) 2372–2388. doi:10.1016/J.JMB.2017.10.018.

[12] R. Giri, D. Kumar, N. Sharma, V.N. Uversky, Intrinsically disordered side of the Zika virus proteome, Front. Cell. Infect. Microbiol. 6 (2016) 144. doi:10.3389/FCIMB.2016.00144/BIBTEX.

[13] W.W. Phoo, Y. Li, Z. Zhang, M.Y. Lee, Y.R. Loh, Y.B. Tan, E.Y. Ng, J. Lescar, C. Kang, D. Luo, Structure of the NS2B-NS3 protease from Zika virus after self-cleavage, Nat. Commun. 2016 71. 7 (2016) 1–8. doi:10.1038/ncomms13410.

[14] X. Chen, K. Yang, C. Wu, C. Chen, C. Hu, O. Buzovetsky, Z. Wang, X. Ji, Y. Xiong, H. Yang, Mechanisms of activation and inhibition of Zika virus NS2B-NS3 protease, Cell Res. 2016 2611. 26 (2016) 1260–1263. doi:10.1038/cr.2016.116.

[15] L. Lim, G. Gupta, A. Roy, J. Kang, S. Srivastava, J. Shi, J. Song, Structurally- and dynamically-driven allostery of the chymotrypsin-like proteases of SARS, Dengue and Zika viruses, Prog. Biophys. Mol. Biol. 143 (2019) 52–66. doi:10.1016/J.PBIOMOLBIO.2018.08.009.

[16] A. Kumar, P. Kumar, R. Giri, Zika virus NS4A cytosolic region (residues 1–48) is an intrinsically disordered domain and folds upon binding to lipids, Virology. 550 (2020) 27–36. doi:10.1016/J.VIROL.2020.07.017.

[17] J. Jumper, R. Evans, A. Pritzel, T. Green, M. Figurnov, O. Ronneberger, K. Tunyasuvunakool, R. Bates, A. Žídek, A. Potapenko, A. Bridgland, C. Meyer, S.A.A. Kohl, A.J. Ballard, A. Cowie, B. Romera-Paredes, S. Nikolov, R. Jain, J. Adler, T. Back, S. Petersen, D. Reiman, E. Clancy, M. Zielinski, M. Steinegger, M. Pacholska, T. Berghammer, S. Bodenstein, D. Silver, O. Vinyals, A.W. Senior, K. Kavukcuoglu, P. Kohli, D. Hassabis, Highly accurate protein structure prediction with AlphaFold, Nat. 2021 5967873. 596 (2021) 583–589. doi:10.1038/s41586-021-03819-2.

[18] G. Madhavi Sastry, M. Adzhigirey, T. Day, R. Annabhimoju, W. Sherman, Protein and ligand preparation: parameters, protocols, and influence on virtual screening enrichments, J. Comput. Aided. Mol. Des. 27 (2013) 221–234. doi:10.1007/s10822-013-9644-8.

[19] P. Kumar, K.U. Saumya, R. Giri, Identification of peptidomimetic compounds as potential inhibitors against MurA enzyme of Mycobacterium tuberculosis, J. Biomol. Struct. Dyn. (2019) 1–21. doi:10.1080/07391102.2019.1696231.

[20] W.L. Jorgensen, D.S. Maxwell, J. Tirado-Rives, Development and Testing of the OPLS All-Atom Force Field on Conformational Energetics and Properties of Organic Liquids, J. Am. Chem. Soc. 118 (1996) 11225–11236. doi:10.1021/ja9621760.

[21] D. Shivakumar, E. Harder, W. Damm, R.A. Friesner, W. Sherman, Improving the prediction of absolute solvation free energies using the next generation opls force field, J. Chem. Theory Comput. 8 (2012) 2553–2558. doi:10.1021/ct300203w.

[22] The PRODRG Server, (n.d.). http://davapc1.bioch.dundee.ac.uk/cgi-bin/prodrg (accessed February 20, 2022).

[23] A. Soranno, I. Koenig, M.B. Borgia, H. Hofmann, F. Zosel, D. Nettels, B. Schuler, Single-molecule spectroscopy reveals polymer effects of disordered proteins in crowded environments, Proc. Natl. Acad. Sci. U. S. A. 111 (2014) 4874–4879. doi:10.1073/PNAS.1322611111/-/DCSUPPLEMENTAL.

[24] R.J. Ellis, Macromolecular crowding: obvious but underappreciated, Trends Biochem. Sci. 26 (2001) 597–604. doi:10.1016/S0968-0004(01)01938-7.

[25] L.F.S. Mendes, L.G.M. Basso, P.S. Kumagai, R. Fonseca-Maldonado, A.J. Costa-Filho, Disorder-to-order transitions in the molten globule-like Golgi Reassembly and Stacking Protein, Biochim. Biophys. Acta - Gen. Subj. 1862 (2018) 855–865. doi:10.1016/J.BBAGEN.2018.01.009.

[26] A. Kumar, A. Kumar, P. Kumar, N. Garg, R. Giri, SARS-CoV-2 NSP1 C-terminal (residues 131–180) is an intrinsically disordered region in isolation, Curr. Res. Virol. Sci. 2 (2021) 100007. doi:10.1016/J.CRVIRO.2021.100007.

[27] S.A. Shiryaev, A.Y. Strongin, Structural and functional parameters of the flaviviral protease: a promising antiviral drug target., Future Virol. 5 (2010) 593–606. doi:10.2217/fvl.10.39.

[28] M. Bollati, K. Alvarez, R. Assenberg, C. Baronti, B. Canard, S. Cook, B. Coutard, E. Decroly, X. de Lamballerie, E.A. Gould, G. Grard, J.M. Grimes, R. Hilgenfeld, A.M. Jansson, H. Malet, E.J. Mancini, E. Mastrangelo, A. Mattevi, M. Milani, G. Moureau, J. Neyts, R.J. Owens, J. Ren, B. Selisko, S. Speroni, H. Steuber, D.I. Stuart, T. Unge, M. Bolognesi, Structure and functionality in flavivirus NS-proteins: perspectives for drug design., Antiviral Res. 87 (2010) 125–48. doi:10.1016/j.antiviral.2009.11.009.

[29] H.H. Chang, R.G. Huber, P.J. Bond, Y.H. Grad, D. Camerini, S. Maurer-Stroh, M. Lipsitch, Analyse systématique des similarités protéiques entre le virus Zika et d’autres virus transmis par des arthropodes, Bull. World Health Organ. 95 (2017) 517–525. doi:10.2471/BLT.16.182105.

[30] C.G. Noble, C.C. Seh, A.T. Chao, P.Y. Shi, Ligand-Bound Structures of the Dengue Virus Protease Reveal the Active Conformation, J. Virol. 86 (2012) 438–446. doi:10.1128/JVI.06225-11/ASSET/D720D1A8-7246-43BA-95DB-E259D4A14D6A/ASSETS/GRAPHIC/ZJV9990954710006.JPEG.

[31] G. Gupta, L. Lim, J. Song, NMR and MD Studies Reveal That the Isolated Dengue NS3 Protease Is an Intrinsically Disordered Chymotrypsin Fold Which Absolutely Requests NS2B for Correct Folding and Functional Dynamics, PLoS One. 10 (2015) e0134823. doi:10.1371/journal.pone.0134823.

[32] D. Kumar, P.M. Mishra, K. Gadhave, R. Giri, Conformational dynamics of p53 N-terminal TAD2 region under different solvent conditions, Arch. Biochem. Biophys. 689 (2020) 108459. doi:10.1016/J.ABB.2020.108459.

[33] A. Kumar, P. Kumar, S. Kumari, V.N. Uversky, R. Giri, Folding and structural polymorphism of p53 C-terminal domain: One peptide with many conformations, Arch. Biochem. Biophys. 684 (2020) 108342. doi:10.1016/J.ABB.2020.108342.

