## Supplemental File for "Investigating the folding dynamics of NS2B protein of Zika virus"

**Supplementary information**

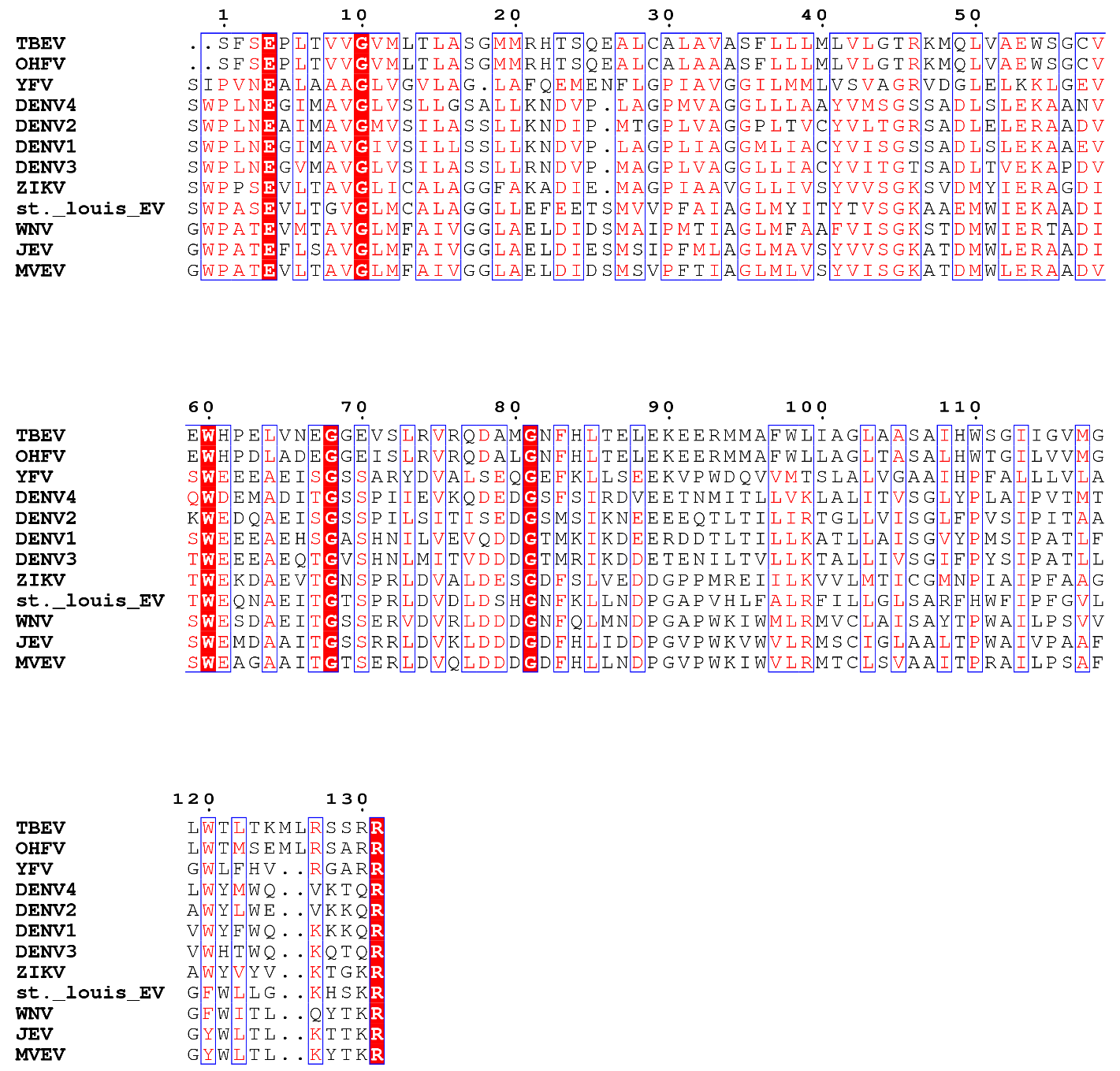

**Figure S1.** **Multiple sequence alignment of NS2B full-length protein of flaviviruses using clustal omega.** The amino acid sequence of NS2B protein for the respective flaviviruses was taken from UniProt database of protein sequence (ZIKV- Brazilian isolate BeH823339; DENV1- strain Nauru/West Pac/1947, P17763; DENV2-strain Thailand/16681/1984, P29990; DENV3-strain Sri Lanka/1266/2000, Q6YMS4; DENV4-strain Singapore/8976/1995, Q5UCB8; YFV-strain Ghana/Asibi/1927, Q6DV88; WNV-strain NY-99, Q9Q6P4; JEV-strain SA-14, P27395; TBEV-strain Hypr, Q01299; MVEV- strain MVE-1-51, P05769; OHFV- Q7T6D2; St. Louis encephalitis virus(St. Louis EV)-strain MS1-7, P09732).

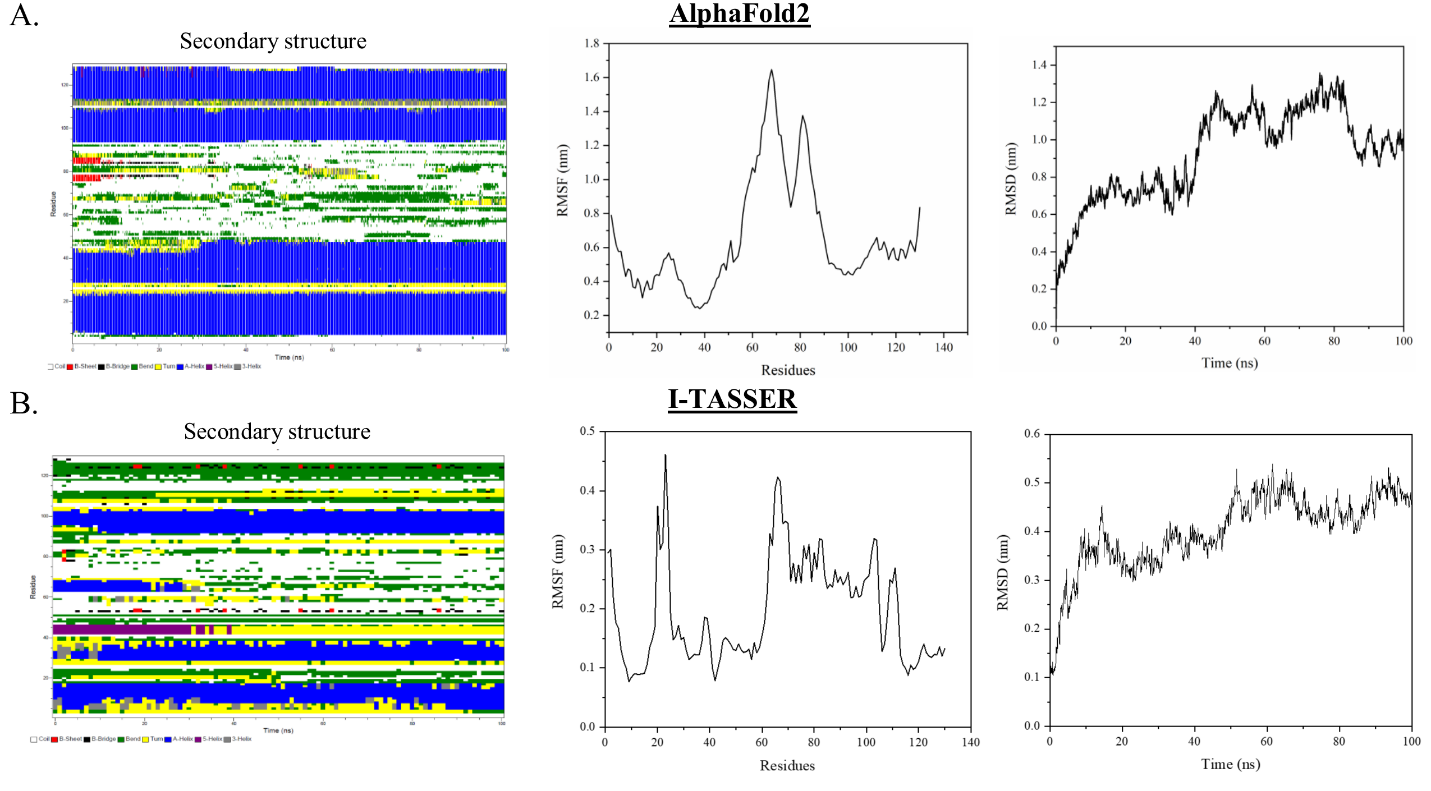

**Figure S2. Parameters associated with ZIKV NS2B full-length protein during MD simulation.** (A) MD simulation parameter of NS2B full-length protein modeled by AlphaFold2 modeling system. 1st panel shows the residue's contribution to the secondary structure of NS2B. The Blue colored segment indicates the presence of 4 helical regions (5-24, 29-48, 94-109, & 114-128). 2^nd^ and 3^rd^ panels show the RMSF of each residue and RMSD during the simulation period. (B) MD simulation parameter of NS2B full-length protein modeled by I-TASSER modeling system. 1^st^ panel shows the residue's contribution to the secondary structure of NS2B. The Blue colored segment in this panel indicates the presence of 4 helical regions (1-20, 30-40, 64-67 & 115-125). 2^nd^ and 3^rd^ panels show the RMSF of each residue and RMSD during the simulation period.

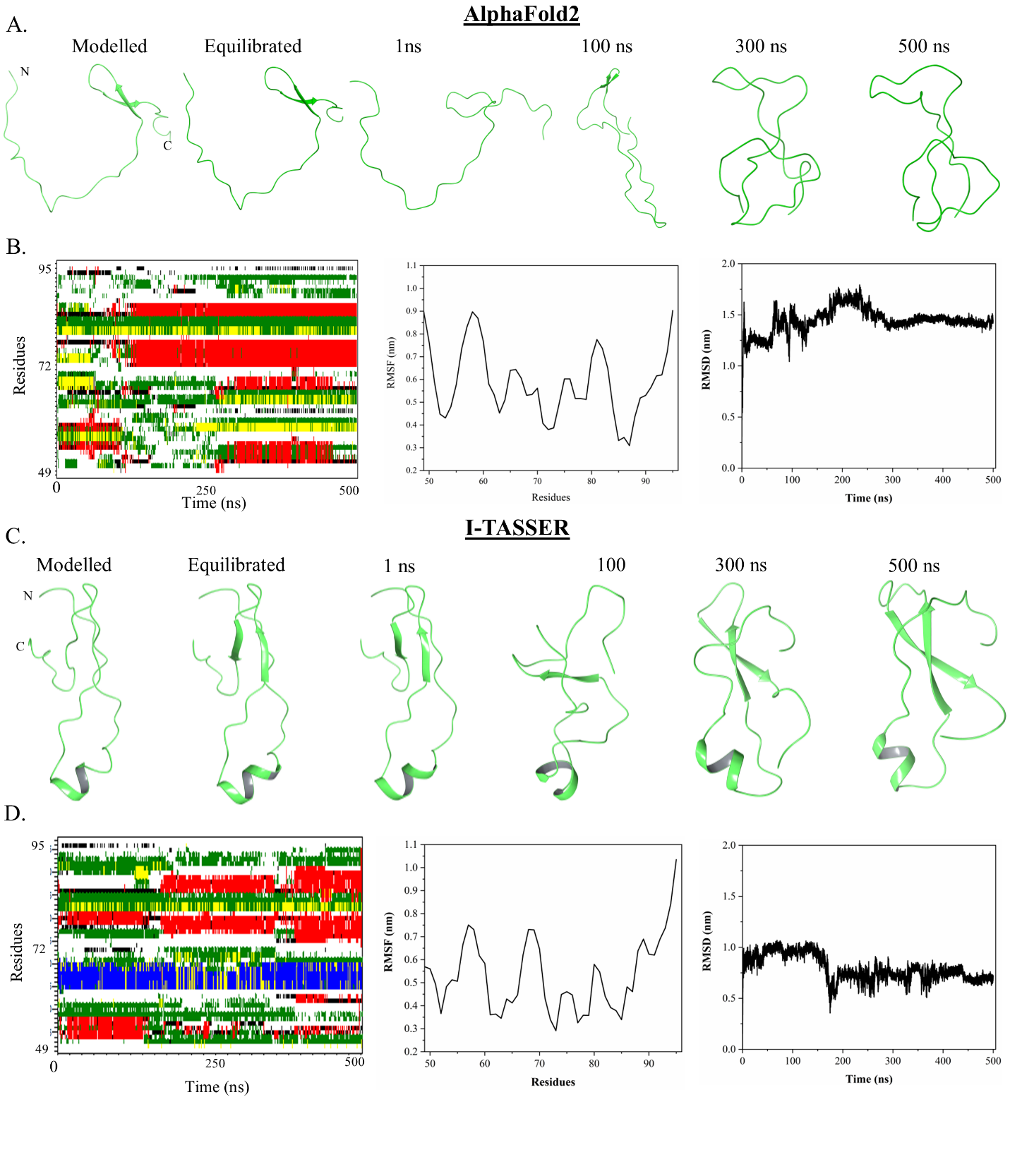

**Figure S3. MD simulation of ZIKV NS2B cytosolic domain (residues 49-95) upto 500 ns.** (A.) Snapshot of the different time frames of NS2B (49-95) during MD simulation in water. The ranked_0 modeled structure predicted by AlphaFold2 modeling system was taken as a template structure for simulation. (B) The first panel shows the residue's contribution to the secondary structure of NS2B (residues 49-95). The Blue colored segment indicates the presence of the helix, and the red-colored segment indicates the strand. 2^nd^ and 3^rd^ panels show the RMSF of each residue and RMSD during the simulation. (C) Snapshot of the different time frames of NS2B (49-95) during MD simulation in water. The model structure predicted by I-TASSER modeling system with a C-score of -1.32 was taken as a template structure for MD simulation. (D) 1^st^ panel shows the residues' contribution to the secondary structure of NS2B (49-95). The Blue colored segment indicates the presence of the helix, and the red-colored segment indicates the strand. 2^nd^ and 3^rd^ panels show the RMSF of each residue and RMSD during the simulation.
